# Cause-excluded Effect Evaluation: a framework for the faithful assessment of genetic perturbation target identification

**DOI:** 10.64898/2026.10.03.752923

**Authors:** Jiaxin Hu, Xiaoning Qi, Limeng Tian, Jie Yan, Nian Liu, Xin Li

**Author notes:** Contributing authors. These authors contributed equally to this work (Jiaxin Hu, Xiaoning Qi).

## Abstract

Identifying which gene drives a desired cell-state transition is a foundational inverse problem in target discovery and cell-state engineering, yet far less developed than forward prediction. We show that prevailing benchmarks inflate performance through two artefacts: training leakage, where test-perturbation cells are seen during fitting, and evaluation leakage, where the perturbed gene’s own expression change exposes the answer. Here, we introduce Cause-excluded Effect Evaluation (CEE), a framework that separates train and test perturbation targets, removes target-gene self-effects from expression profiles and scores candidates in downstream response space. With CEE, we find that simple structure-based methods consistently outperform complex deep learning models across 16 single-cell perturbation datasets, revealing that the recoverable signal is governed by low-rank and additive structure in perturbation responses rather than by model complexity. Our work disentangles evaluation artefacts from genuine predictive capacity and offers a design principle for developing perturbation target identification methods.

## 1 Introduction

Genome-scale perturbation screens coupled to single-cell transcriptomics now make it possible to map genetic interventions to transcriptional state changes across thousands of genes and cells ^1–3^. Currently, most computational methods are designed for forward prediction, in which a model receives a cell state and a specified perturbation and predicts the resulting transcriptional response. Yet many consequential biological design problems require the inverse direction, such as target discovery, phenotypic screen deconvolution, drug repurposing and cell-state engineering. In these scenarios, investigators usually begin from a desired transcriptional state and need to find which gene should be perturbed to reach a goal cell state ^4–8^.

To accomplish perturbation target identification, current methods follow several strategic routes. Some models were designed natively for target identification, such as CausalDN ^9^, PDGrapher ^10^ and CellNavi ^11^. Others repurposed forward prediction models for the inverse task, such as GEARS ^12^, scGPT ^13^ and CPA ^14^, which score candidate perturbations against an observed target state. Public benchmarks typically compare these approaches with differential gene expression (DGE) ranking, which prioritizes genes based on their observed expression change between perturbed and control cells. In this setting, benchmark performance may reflect ranking ability more than recovery of actionable targets from downstream biological response.

Real applications, however, require recovery of actionable perturbation targets from downstream biological response, not merely gene ranking. Whether current benchmarks could measure this biological capability remains to be investigated. Investigating these approaches, we identified two leakage issues through which a method can appear to perform well without recovering meaningful targets from the biologically relevant downstream response, as opposed to merely ranking genes by their direct self-effect. The first, training leakage, occurs when cells carrying test perturbations are used during model fitting, allowing a model to memorize test signatures rather than generalize to unseen targets ^15^. The second, evaluation leakage, arises because DGE ranking is often dominated by the direct expression change of the target gene itself in the query profile rather than by downstream transcriptional consequences, so that scoring can conflate this self-effect signal with genuine downstream response. Related benchmarks for forward prediction have raised parallel concerns about systematic variation inflating apparent generalization ^16,17^. While these issues motivate the need for improved evaluation, self-effect leakage remains specific to the inverse task and has not been addressed by those efforts. It arises from direct self-suppression of the knocked-down gene in CRISPR interference (CRISPRi) screens, a phenomenon absent from forward prediction. A benchmark that excludes this self-effect information while preserving downstream responses has therefore been lacking.

Here, we introduce Cause-excluded Effect Evaluation (CEE), a benchmarking framework that addresses both training and evaluation leakage. CEE evaluation separates train and test perturbation targets using a perturbation-level out-of-distribution split, excludes all cells from each held-out target gene from training, removes target-gene self-effects at the expression-profile level, and scores candidate perturbations in the full downstream response space. We apply this framework to 16 single-cell perturbation datasets spanning 12 single-gene and 4 dual-gene settings, and benchmark statistical baselines, native target-identification models and repurposed forward predictors. Under CEE evaluation, simple structure-based methods consistently outperform complex deep learning models, whereas standard and upper-bound evaluation settings overstate performance. We further show that low-rank and additive structure in the perturbation response landscape explains much of the recoverable signal, defining a learnability boundary for target identification methods. These findings complement recent forward-prediction benchmarks that identified similar gaps between deep learning complexity and task-appropriate signal ^16,17^, and extend them to the underexplored inverse direction.

## 2 Results

### 2.1 CEE separates perturbation target identity from downstream transcriptional response to enable faithful benchmarking

We introduce CEE, a benchmarking framework that formalizes perturbation target identification as a retrieval problem over genetic targets and addresses two distinct failure modes in existing evaluation protocols. In Perturb-seq experiments ^1^, a perturbation yields a differential expression signature Δ*x* = *x*_pert_ *− x*_ctrl_, and the inverse task asks whether a method can infer the perturbed target gene *g^∗^* from this signature (Fig. 1a). Faithful evaluation requires that *g^∗^* be recoverable from the downstream functional response captured in Δ*x*, not from artefacts that directly encode the answer. Training leakage arises when test-perturbation cells are observed during model fitting (Fig. 1b). Evaluation leakage arises when the target gene’s own self-effect dominates the query signature, exposing *g^∗^* without requiring any modeling of downstream biology (Fig. 1c).

**Fig. 1.**
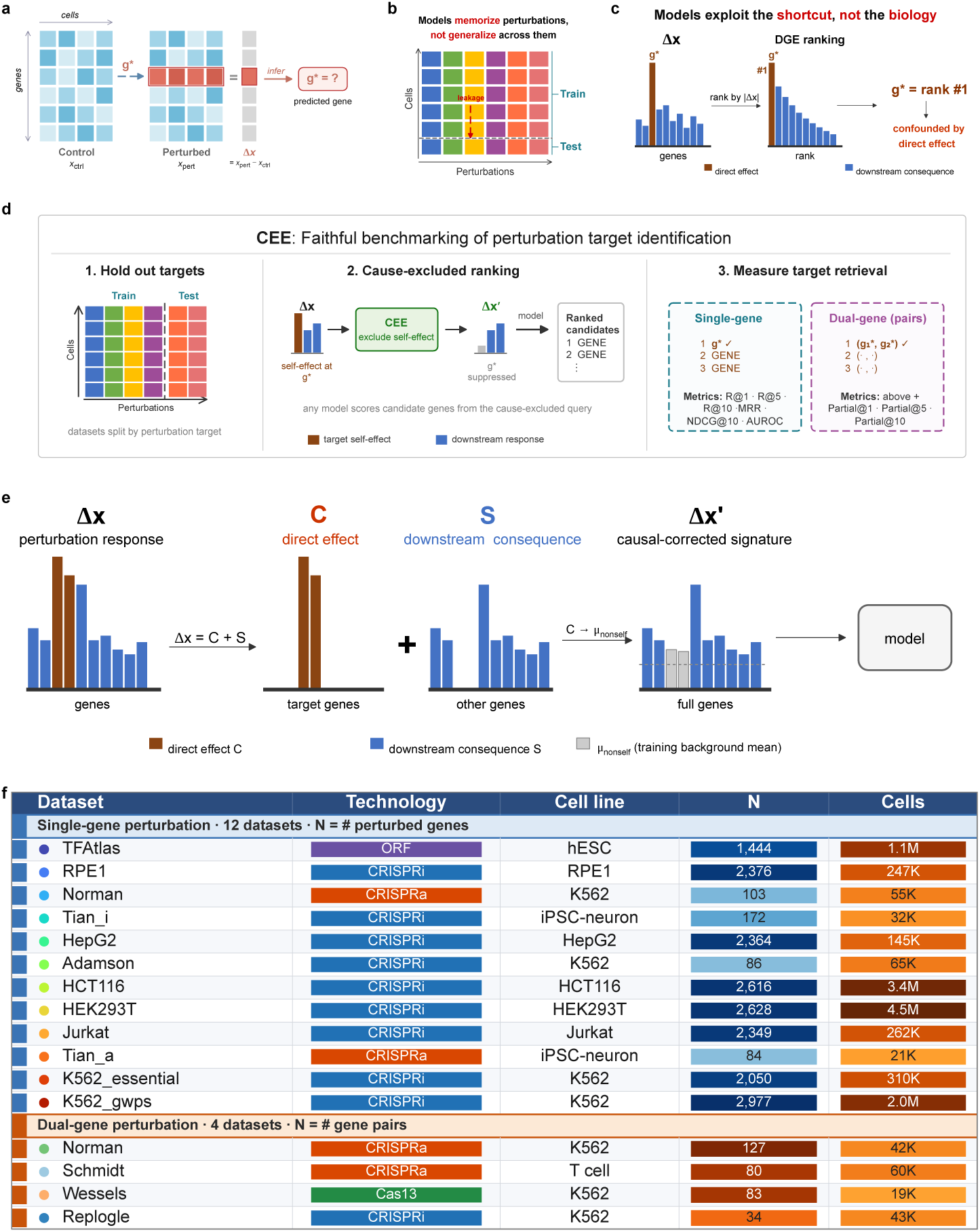
Perturbation target identification, sources of benchmark leakage, and the Cause-excluded Effect Evaluation (CEE) framework. a, The perturbation target identification task. A perturbation of target gene *g^∗^* transforms control cells *x*_ctrl_ into a perturbed state *x*_pert_, and the differential expression signature Δ*x* = *x*_pert_ *− x*_ctrl_ serves as the query from which *g^∗^* is inferred. **b**, Training leakage under a cell-level data split. Each column is a perturbation and each row is a cell, and a dashed line divides the cells of every perturbation into training and test partitions. Because cells from the same perturbations populate both partitions, a model can recognize at test time the perturbations it has already encountered during fitting, memorizing perturbation identity rather than generalizing across perturbations. **c**, Evaluation leakage from the direct self-effect of the target gene. The observed response Δ*x* carries a large direct effect at the target gene *g^∗^* alongside smaller downstream signals, so ranking genes by *|*Δ*x|* places *g^∗^* at rank 1. **d**, The CEE evaluation workflow. Each dataset is first split by perturbation target, and all cells of the held-out target genes are excluded from model fitting (step 1). The cause-excluded query is then scored by any candidate model, which ranks genes by their compatibility with the downstream response (step 2). Target retrieval is finally assessed for single-gene and dual-gene perturbations using rank-based metrics, including recall at rank *k* (R@*k*), mean reciprocal rank (MRR), normalized discounted cumulative gain (NDCG@10) and area under the receiver operating characteristic curve (AUROC), with partial recall (Partial@*k*) added for gene pairs (step 3). **e**, The CEE profile-wise self-effect exclusion procedure. The response is decomposed as Δ*x* = *C* + *S* into a target-gene direct-effect component *C* and a downstream-consequence component *S*. The target-gene dimensions are replaced by the non-self background mean *µ*_nonself_ estimated from training-set perturbations, yielding the CEE-corrected signature Δ*x^′^* that is passed to the model. **f**, The 16 benchmark datasets, comprising 12 single-gene and 4 dual-gene perturbation collections that span CRISPR interference (CRISPRi), CRISPR activation (CRISPRa), open reading frame (ORF) and Cas13 modalities. *N* denotes the number of perturbed genes for single-gene datasets or gene pairs for dual-gene datasets, and Cells denotes the number of single-cell transcriptomes profiled.

CEE resolves training leakage by adopting a perturbation-level out-of-distribution split in which all cells from each held-out target gene are excluded from model fitting. CEE resolves evaluation leakage by decomposing each perturbation response into target-gene direct-effect dimensions *C* and downstream-consequence dimensions *S*, then replacing the target-gene dimensions with a fold-specific non-self background mean estimated from training perturbations. This produces a CEE-corrected signature Δ*x^′^* in which the direct target-gene self-effect is suppressed while the downstream expression outcome is retained (Fig. 1d,e).

We applied this framework to 16 single-cell perturbation datasets comprising 12 single-gene and 4 dual-gene datasets, spanning CRISPRi, CRISPR activation (CRISPRa), open reading frame (ORF) and Cas13 technologies across multiple cell lines ^3,18–26^ (Fig. 1f). This collection enables systematic evaluation of how each leakage mode shapes apparent method performance across diverse experimental contexts.

### 2.2 Training and evaluation leakage jointly inflate performance estimates in current single-gene benchmarks

We examined how training leakage and evaluation leakage each contribute to current single-gene benchmark outcomes. Under the standard evaluation setting, CausalDN, PDGrapher and CellNavi ^9–11^ produced high recall at rank 1 (R@1) values across multiple datasets, with CellNavi ranking first across all summary metrics. DGE ranks candidates by differential expression magnitude and is retained as an upper-bound reference in standard evaluation, but is excluded from CEE fair ranking. Its performance increased with the self-effect ratio *r* = *|*Δ_obs_[*g^∗^*]*|*/ max*_g_ |*Δ_obs_[*g*]*|*, which measures how strongly the target gene dominates the observed expression profile. On TFAtlas ^21^, where ORF overexpression produces a trivial perturbation fraction of only 0.1% and rarely places the target gene at the top of the differential expression ranking, all methods including DGE remained near the performance floor (Fig. 2a). Current benchmarks thus produce an optimistic performance landscape that depends critically on how strongly the target-gene self-effect is represented in the query profile.

**Fig. 2.**
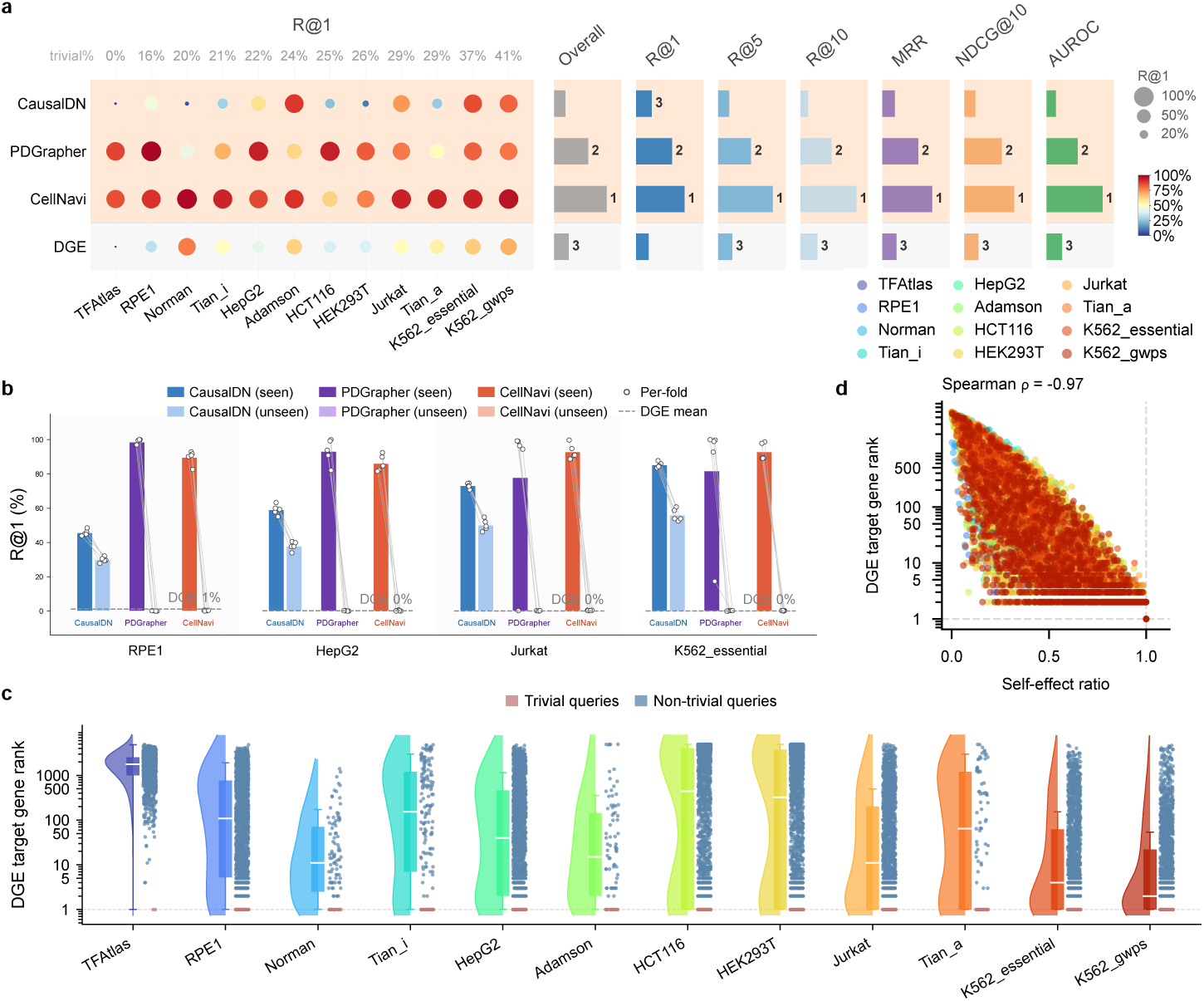
Current single-gene benchmarks expose training and evaluation leakage. **a**, Recall at rank 1 (R@1) bubble matrix for CausalDN, PDGrapher and CellNavi under the current evaluation setting, with differential gene expression (DGE) as a reference, across 12 single-gene datasets. Bubble area and color encode R@1 for each method and dataset. Datasets are ordered by increasing fraction of trivial queries, annotated above the matrix. The right panels show normalized rank scores for Overall, R@1, R@5, R@10, mean reciprocal rank (MRR), normalized discounted cumulative gain at rank 10 (NDCG@10) and area under the receiver operating characteristic curve (AUROC), with the top three methods labeled for each metric. **b**, R@1 for CausalDN, PDGrapher and CellNavi on RPE1, HepG2, Jurkat and K562_essential under the seen and unseen perturbation splits. Bars show the mean across five cross-validation folds, points show per-fold values, grey lines connect paired folds, and the dashed line marks the DGE mean for each dataset. **c**, Distribution of DGE target-gene ranks across 12 single-gene datasets, shown on a log scale. Each half-violin is a kernel density estimate; the box marks the median and interquartile range, and whiskers extend to 1.5 times the interquartile range. Points denote trivial and non-trivial queries, and the dashed line marks rank 1. **d**, Per-query self-effect ratio versus DGE target-gene rank across 12 single-gene datasets, with the self-effect ratio defined as *|*Δ_obs_[*g^∗^*]*|*/ max*_g_ |*Δ_obs_[*g*]*|* for target gene *g^∗^*. Colours identify datasets, dashed lines mark ratio 1 and rank 1, and the Spearman correlation is *ρ* = *−*0.97.

To isolate training leakage, we evaluated CausalDN, PDGrapher and CellNavi under an unseen-perturbation split that masks test-perturbation cells during training while preserving the target-gene self-effect in the evaluation signature. Across RPE1, HepG2, Jurkat and K562_essential ^3,19^, moving from seen to unseen perturbations substantially reduced R@1 across all three methods (Fig. 2b). CausalDN dropped by 15–29 percentage points, whereas PDGrapher and CellNavi dropped by 78–99 percentage points, indicating that seen-split performance derived largely from memorizing test signatures.

We then examined evaluation leakage through the DGE target-gene rank distribution. A perturbation is defined as trivial when the target gene *g^∗^* has the largest absolute expression change in Δ_obs_, so that DGE directly exposes *g^∗^* at rank 1. Across all 12 single-gene datasets, trivial queries concentrated tightly at rank 1 under DGE scoring, whereas non-trivial queries produced broadly distributed and generally higher ranks (Fig. 2c).

To quantify this dependence continuously, we computed *r* for each query and measured its association with the DGE target-gene rank across all 12 datasets. The self-effect ratio was strongly and inversely correlated with rank (Spearman *ρ* = *−*0.97, Fig. 2d). The relative magnitude of the target gene’s expression change accounted for nearly all variance in DGE rankings across technologies and cell lines. This implies that any benchmark relying on DGE as a reference or upper bound is not measuring target identification ability but rather the exploitability of the self-effect signal. Faithfully evaluating downstream-response-based retrieval therefore requires removing the target-gene self-effect from the evaluation signature before ranking.

### 2.3 Self-effect signals drive evaluation leakage and CEE correction preserves downstream response energy

We further dissected the mechanism by which target-gene self-effects enter differential-expression ranking. The self-effect signal refers to the direct expression change of the target gene *g^∗^* itself in Δ_obs_, distinct from downstream regulatory consequences. To isolate this contribution experimentally, we introduced a controlled perturbation modification in K562_gwps ^3^. We selected three perturbations whose target genes showed absolute log fold-change values below 1.0. We replaced each target-gene dimension with the per-fold control mean and simultaneously downscaled *HSP90AB1* to 0.2*×* its control level, injecting an absolute log fold-change of approximately 1.0 that exceeded all three true target-gene changes. The overall UMAP organization of perturbed and control cells was largely unchanged (Fig. 3a), but the top DGE-ranked gene switched from the true target to *HSP90AB1* for all three perturbations (Fig. 3b). This counterfactual experiment demonstrates that DGE identifies whichever gene carries the largest expression change in the query profile, not the perturbation target from downstream biology.

**Fig. 3.**
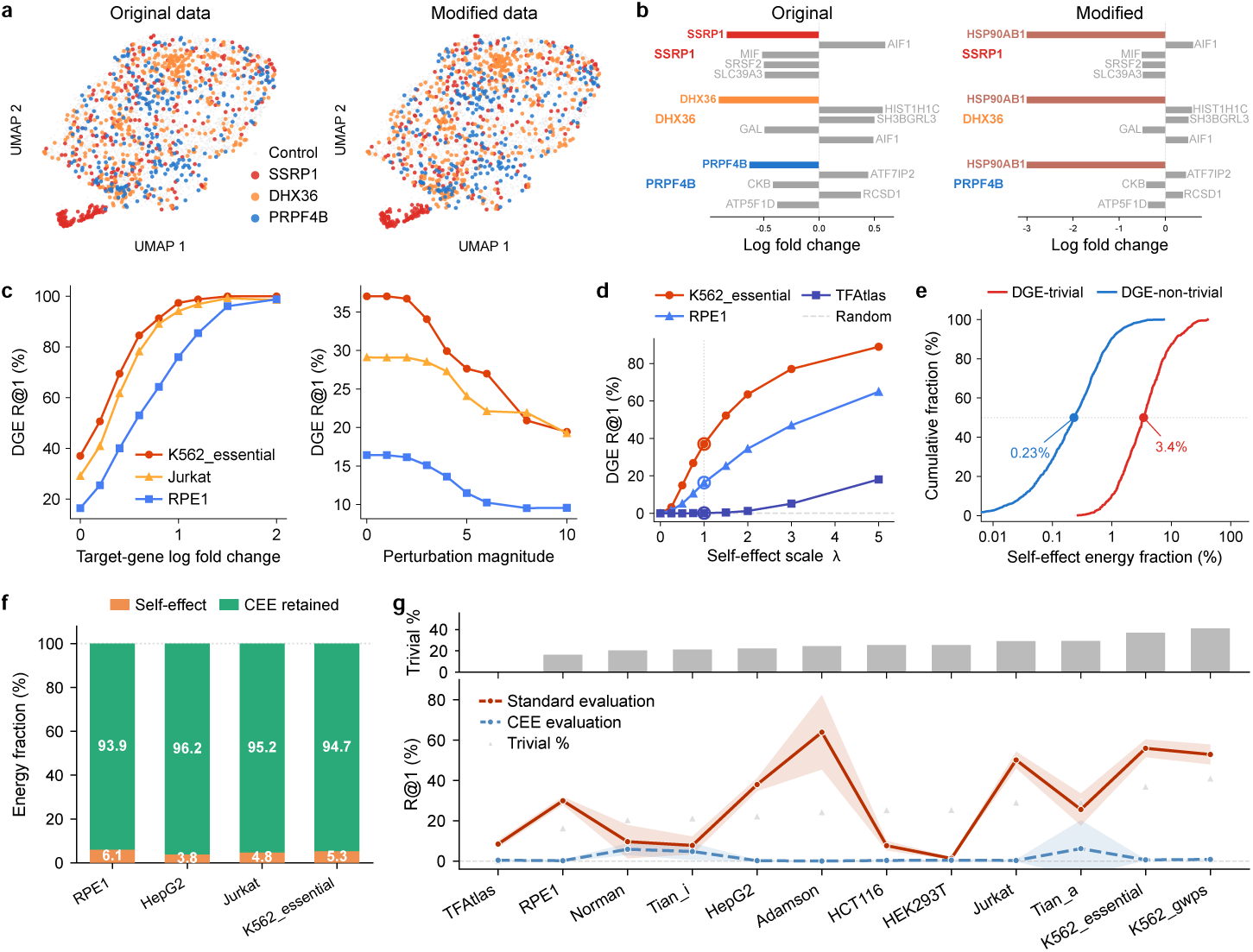
Self-effect-driven differential gene expression (DGE) behavior and Cause-excluded Effect Evaluation (CEE) evaluation. **a**, UMAP embeddings of K562_gwps cells before and after expression modification. Colours identify control cells and the *SSRP1*, *DHX36* and *PRPF4B* perturbation populations. **b**, Top differentially expressed genes by log fold change for the same three perturbations before and after modification, in which each target-gene dimension is set to the control mean and *HSP90AB1* is downscaled in perturbed cells. In the original data each target gene ranks first, whereas after modification *HSP90AB1* ranks first for all three perturbations. **c**, DGE recall at rank 1 (R@1) as a function of the target-gene log fold-change threshold (left) and the perturbation-magnitude threshold (right) in K562_essential, Jurkat and RPE1. Each point represents the subset of perturbations passing the corresponding threshold. **d**, DGE R@1 after scaling the target-gene self-effect by a factor *λ* in K562_essential, RPE1 and TFAtlas. Open circles mark *λ* = 1, corresponding to the unmodified data, and the dashed line marks the random baseline. **e**, Empirical cumulative distributions of the self-effect energy fraction for DGE-trivial and DGE-nontrivial perturbations in K562_essential. Filled points mark the group medians of 3.4% and 0.23%, respectively. **f**, Self-effect and CEE-retained energy fractions for DGE-trivial perturbations across four CRISPR interference (CRISPRi) datasets, with each bar partitioned into a self-effect fraction and a CEE-retained fraction. **g**, CausalDN-Synthetic R@1 across 12 single-gene datasets under standard and CEE evaluation. The upper bar plot shows the fraction of trivial queries per dataset, and the lower plot shows the mean R@1 across five folds with shaded 95% confidence intervals; grey triangles mark the trivial-query fraction.

We next quantified self-effect dependence through two continuous dimensions. In K562_essential, Jurkat and RPE1^3,19^, DGE R@1 increased monotonically as the target-gene log fold-change threshold was raised, and decreased monotonically as the perturbation-magnitude threshold increased (Fig. 3c). To establish causality, we scaled the target-gene self-effect by a multiplicative factor *λ* from 0 to 5.0 in K562_essential, RPE1 and TFAtlas ^21^. DGE R@1 increased monotonically with *λ* in all three datasets, and at *λ* = 0 performance dropped to approximately 0% (Fig. 3d). TFAtlas uses ORF overexpression with a natural trivial fraction of 0.1%, yet showed the same monotonic response, ruling out the possibility that self-effect dependence is a CRISPRi-specific artifact.

We then evaluated whether CEE exclusion removes a biologically meaningful fraction of the perturbation response. In K562_essential, the median self-effect energy fraction was 3.4% for trivial perturbations and 0.23% for non-trivial perturbations (Fig. 3e). Across four CRISPRi datasets spanning RPE1, HepG2, Jurkat and K562_essential, CEE exclusion removed 3–6% of perturbation response energy and retained 94–97% in the downstream-consequence dimensions (Fig. 3f). CEE thus acts as a targeted correction on a small fraction of the expression profile rather than a broad removal of perturbation signal. This small energy removal translated into a 48.7 percentage-point reduction in DGE R@1, revealing that rank-based metrics are disproportionately sensitive to which dimensions carry the signal. The self-effect dimensions are energetically minor but informationally decisive for DGE. CEE exclusion is therefore a prerequisite for fair evaluation rather than a form of information loss.

Finally, we tested whether self-effect dependence extends to model-based inverse predictors. We evaluated CausalDN-Synthetic, a zero-shot predictor that uses synthetic perturbation representations, under both standard and CEE evaluation across 12 single-gene datasets. Under standard evaluation, its R@1 increased with the trivial perturbation fraction (*ρ ≈ −*0.97). Under CEE evaluation, R@1 was reduced to near-zero levels across all datasets (Fig. 3g). Self-effect leakage therefore operates at the level of the evaluation protocol and affects model-based methods as well as ranking baselines.

### 2.4 CEE evaluation reveals limited but biologically structured learnability in single-gene target identification

We performed a comprehensive single-gene benchmark under CEE evaluation, comparing 12 fair-ranking methods across six groups: native reverse models (CausalDN ^9^, PDGrapher ^10^, CellNavi ^11^), forward models repurposed for retrieval (GEARS ^12^, scGPT ^13^, CPA ^14^), response-imputation approaches (SVD-linear, k-NN), gene-feature methods (Coexpr, Gene2Sig) and reverse classifiers. DGE was retained as an upper-bound reference (Fig. 4a). All 12 fair methods showed R@1 below 3.1%, with SVD-linear and k-NN achieving the highest values. The DGE upper-bound reference reached 48.7%, more than 15-fold above the best fair method (Fig. 4b). This gap does not reflect a difference in biological signal quality but a difference in what is being measured. Under leaky conditions, the top-performing methods were those that best exploited the self-effect shortcut. Under CEE evaluation, that shortcut was absent, and the same methods reverted to near-random performance. The existing benchmark record therefore describes how well methods leverage self-effect information, not how well they identify targets from downstream biological response.

**Fig. 4.**
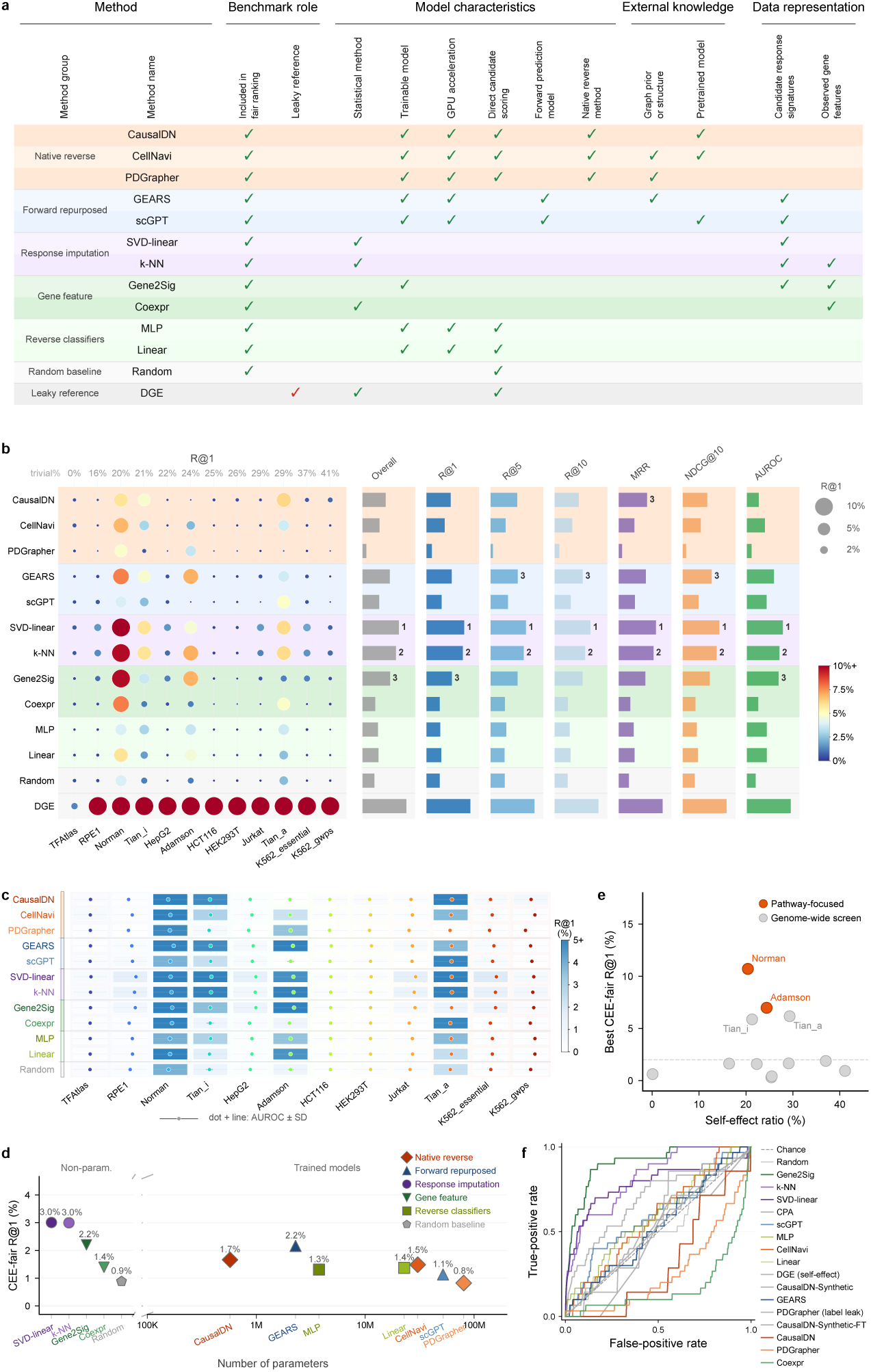
Single-gene perturbation target identification benchmark under Cause-excluded Effect Evaluation (CEE). **a**, Attribute matrix for the 13 methods assessed in the single-gene benchmark. The 12 fair-ranking methods are organized into six groups (Native reverse, Forward repurposed, Response imputation, Gene feature, Reverse classifiers and Random baseline), with differential gene expression (DGE) retained as an upper-bound reference. Columns report binary attributes across four categories: Benchmark role, Model characteristics, External knowledge and Data representation, and check marks indicate the presence of an attribute. **b**, CEE recall at rank 1 (R@1) bubble matrix for all 13 methods across 12 single-gene datasets. Bubble area and color encode R@1 for each method and dataset. Datasets are ordered by increasing fraction of trivial queries, annotated above the matrix, and method groups share the background colors in a, with DGE shown as an upper-bound reference. The right panels show normalized rank scores for Overall, R@1, R@5, R@10, mean reciprocal rank (MRR), normalized discounted cumulative gain at rank 10 (NDCG@10) and area under the receiver operating characteristic curve (AUROC), with the top three methods labeled for each metric. **c**, Dual-metric performance matrix for the 12 fair-ranking methods across 12 single-gene datasets. Cell background color encodes CEE R@1, and within each cell a point and horizontal bar show AUROC *±* s.d. across folds. Methods and datasets are ordered as in b. **d**, Mean CEE R@1 across 12 datasets versus the number of trainable parameters for all 13 methods. Non-parametric methods are shown on the left categorical axis and parametric models on the right log-scaled axis, with marker shape indicating method group and R@1 annotated next to each point. **e**, Best CEE R@1, defined as the maximum across the 12 fair methods, versus self-effect ratio for each single-gene dataset. Pathway-focused datasets (Norman and Adamson) and genome-wide screens are distinguished by color, and the dashed line marks R@1 = 2%. **f**, ROC curves for retrieval of ribosomal protein genes (RPL, RPS, MRPL and MRPS families; *n* = 30) among all perturbed genes in K562_gwps. Each curve is one method, AUC values are listed in the legend, and the diagonal dashed line marks chance (AUC = 0.50).

The low-performance pattern was consistent across complementary metrics. R@1 and area under the receiver operating characteristic curve (AUROC) both converged near chance for most method–dataset pairs, with elevated signal concentrated in Norman ^23^ and Adamson ^18^ (Fig. 4c). Dataset-level variance exceeded method-level variance, indicating that dataset composition was a stronger determinant of performance than method design. No external model class showed meaningful enrichment above chance across the benchmark (Fig. 4b,c).

We also assessed whether model capacity explained the residual performance. SVD-linear and k-NN achieved the highest mean R@1 at 3.0%, whereas PDGrapher at 80M parameters reached only 0.8%, below the random baseline (Fig. 4d). Methods imposing a low-rank or neighborhood prior outperformed those relying on generic parametric capacity or external network priors, indicating that inductive bias rather than model size determined performance under CEE.

To identify the origin of the residual learnable signal, we examined whether the trivial perturbation fraction predicted CEE performance. Dataset-level learnability was uncorrelated with the self-effect ratio. K562_gwps ^3^, despite having the highest trivial fraction, showed CEE R@1 indistinguishable from random, whereas Norman and Adamson, whose perturbation panels target coherent biological pathways, yielded the highest learnability (Fig. 4e). In pathway-focused panels, perturbations within the same functional program produce similar downstream signatures, providing the co-occurrence structure that low-rank and neighborhood methods exploit. This means that what current methods learn under CEE is functional module signature similarity, not the global architecture of gene regulatory networks. Single-gene target identification from an unstructured transcriptome-wide screen therefore remains effectively unsolved.

The methods ranking highest under CEE also showed superior recovery of functionally coherent gene groups. In K562_gwps, Gene2Sig recovered ribosomal protein genes with AUROC 0.89, compared with 0.27 for PDGrapher; SVD-linear and k-NN showed similarly high recovery, whereas PPI-prior methods fell near or below chance (Fig. 4f). This contrast is mechanistically informative. A PPI graph prior actively interferes with functional recovery, because protein–protein interaction network topology does not align with the functional-similarity geometry that reverse retrieval requires. Data-driven functional signatures outperform graph-structural priors precisely because they encode the downstream response space directly. Taken together, CEE evaluation reveals that single-gene perturbation target identification from downstream response remains largely unsolved under fair conditions, while also defining what class of inductive bias is needed to make progress.

### 2.5 Additive statistical structure dominates dual-gene target identification under CEE evaluation

We extended CEE evaluation to dual-gene perturbation target identification. The benchmark comprised 12 fair-ranking methods and one DGE-pair upper-bound reference, spanning external deep learning models, additive statistical methods, simple baselines and random ranking (Fig. 5a). Across four dual-gene datasets, CEE evaluation produced a clear separation between additive methods and external deep learning models. Ridge-kNN and Ridge ranked first and second with mean R@1 of 44.2% and 42.8% across Norman ^23^, Schmidt ^24^ and Wessels ^25^. All six external deep learning models remained below 10% R@1 regardless of model capacity; GEARS ^12^ was the highest at 5.9% (Fig. 5b). Unlike single-gene responses, combinatorial perturbation outcomes are largely decomposable into the sum of individual components, a property that additive methods exploit directly.

**Fig. 5.**
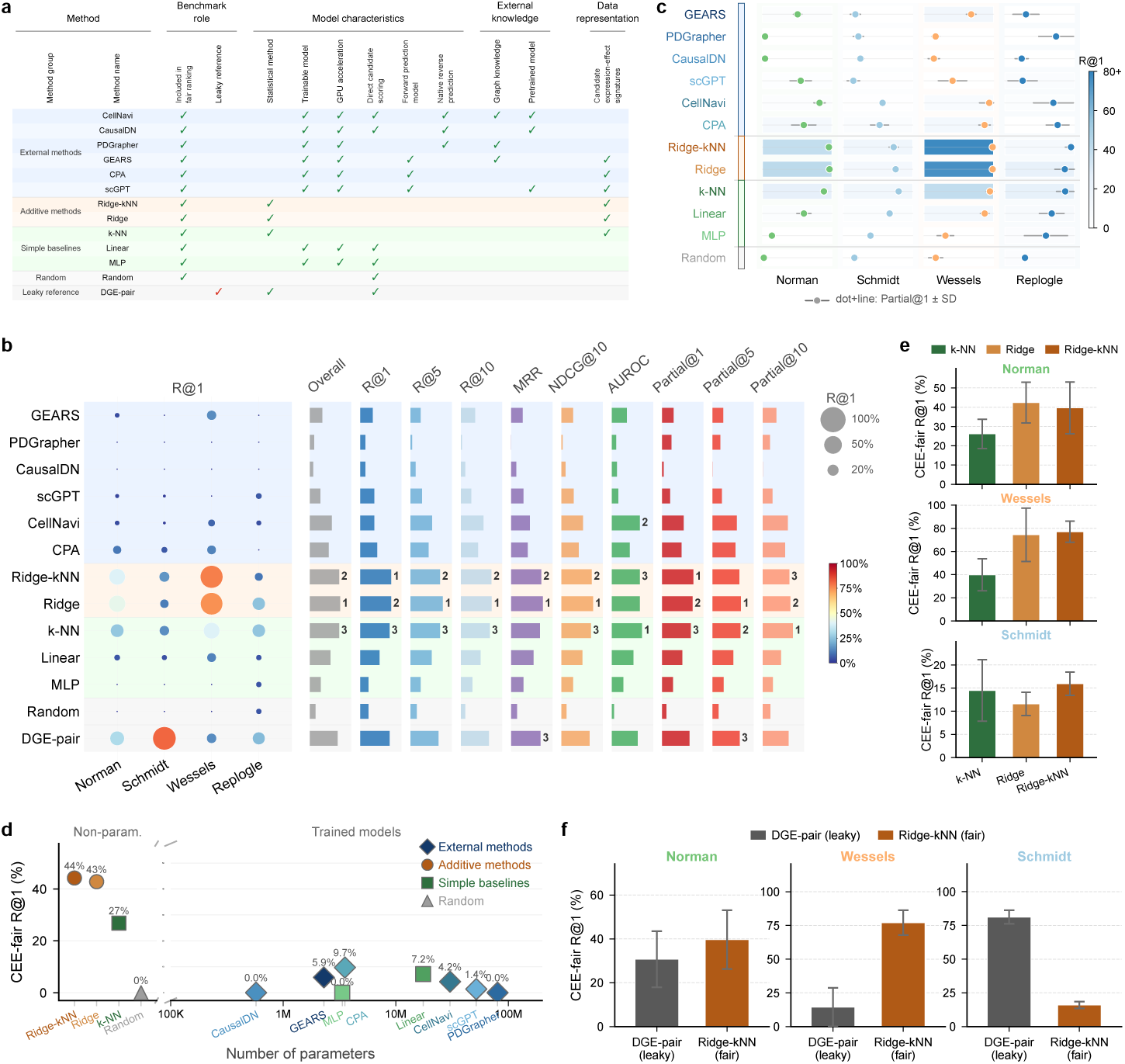
Dual-gene perturbation target identification benchmark under Cause-excluded Effect Evaluation (CEE). **a**, Attribute matrix for the 13 methods assessed in the dual-gene benchmark. The 12 fair-ranking methods are grouped as External methods, Additive methods, Simple baselines and Random, with differential gene expression (DGE) applied to gene pairs (DGE-pair) retained as an upper-bound reference. Columns report binary attributes across Benchmark role, Model characteristics, External knowledge and Data representation; green check marks denote fair-method attributes and red check marks denote the upper-bound reference. **b**, CEE recall at rank 1 (R@1) bubble matrix for the 13 methods across four dual-gene datasets. Bubble area and color encode R@1 for each method and dataset, and grey crosses mark unavailable entries. The right panels show normalized rank scores for Overall, R@1, R@5, R@10, mean reciprocal rank (MRR), normalized discounted cumulative gain at rank 10 (NDCG@10), area under the receiver operating characteristic curve (AUROC), Partial@1, Partial@5 and Partial@10 (partial recall at rank *k*, counting a candidate that shares at least one gene with the true pair), with the top three methods labeled for each metric. **c**, Dual-metric performance matrix for the 12 fair-ranking methods across four dual-gene datasets. Cell background color encodes CEE R@1, and within each cell a point and horizontal bar show Partial@1 *±* s.d. across folds. Methods are grouped as in a and b. **d**, Mean CEE R@1 across Norman, Schmidt and Wessels versus the number of trainable parameters. Non-parametric methods are shown on the left categorical axis and trained models on the right log-scaled axis, with marker shape indicating method group and values annotated next to points. **e**, Contributions of the additive (Ridge) and nearest-neighbor (k-NN) components on Norman, Wessels and Schmidt. For each dataset, bars show CEE R@1 for k-NN, Ridge and Ridge-kNN, with error bars indicating fold s.d. **f**, Per-dataset R@1 for DGE-pair (upper-bound reference) and Ridge-kNN (CEE) on Norman, Wessels and Schmidt. Paired bars show R@1, with error bars indicating fold s.d.

The dual-metric matrix, which jointly displays R@1 and Partial@1 (the fraction of top-ranked candidates sharing at least one gene with the true pair) across methods and datasets, confirmed this advantage at the dataset level. Additive methods maintained their lead across Norman and Wessels, while all external deep learning models remained below 15% R@1 in both datasets (Fig. 5c). Schmidt showed uniformly lower R@1 across all methods due to its small perturbation panel and strong CRISPRa activation signal.

Parameter count did not explain performance in the dual-gene setting. Ridge-kNN and Ridge, which ranked highest overall, achieved these results with no task-specific learned weights. The highest-performing parameterized model, CPA, reached 9.7%, while the largest model, PDGrapher, performed at chance (Fig. 5d). The negative correlation between parameter count and CEE R@1 indicates that the recoverable signal was captured by the additive decomposition assumption rather than model capacity. In this setting, model complexity introduces unconstrained degrees of freedom and functions as a liability rather than a resource, inverting the standard expectation.

To characterize the contribution of each component in Ridge-kNN, we decomposed it into global additive and local neighborhood parts. Moving from k-NN alone to Ridge improved R@1 by up to 34.6 percentage points in large-panel datasets, indicating a dominant contribution from global additive modeling. Schmidt was the exception where k-NN marginally outperformed Ridge, because the smaller candidate set makes local neighborhood information more useful. Ridge-kNN combined both components and produced the most robust cross-dataset behavior among all methods, achieving 15.9% R@1 in Schmidt (Fig. 5e).

Finally, Ridge-kNN exceeded DGE-pair modestly in Norman but by 62.7 percentage points in Wessels. Schmidt was the exception where DGE-pair reached 81.2% compared to Ridge-kNN at 15.9%, reflecting the strong CRISPRa-driven target-gene activation signal in that dataset (Fig. 5f). This exception is itself informative. When the activated target gene dominates the query signature, DGE-pair’s self-effect advantage becomes decisive, confirming that CEE targets precisely this class of signal contamination. Together, these results indicate that dual-gene target identification under CEE evaluation is most reliably solved by the additive decomposability of combinatorial transcriptional responses. This additive geometric prior, rather than model capacity, is the principal design constraint for methods in this setting.

### 2.6 Low-rank and additive structures define the learnable part of perturbation target identification

We examined why methods imposing structured priors outperformed more complex alternatives under CEE evaluation. SVD-linear improved per-perturbation performance over random ranking across the majority of K562_essential ^3^ perturbations, with ribosomal protein genes (RPL/RPS) and proteasome subunit genes (PSMA/PSMB) enriched among perturbations with the largest improvement (Fig. 6a). We computed the singular value spectrum for five CRISPRi datasets and found that cumulative explained variance reached approximately 80% within the first 10 components in all five (Fig. 6b), indicating that perturbation responses are constrained to a low-dimensional subspace. This low-rank geometry means that reverse retrieval need only distinguish perturbations along a handful of functional axes. Despite this low-rank structure, fair-evaluation R@1 remains below 3%, indicating that current methods are not yet effectively aligned with this geometry; the subspace itself represents the principal under-exploited resource.

**Fig. 6.**
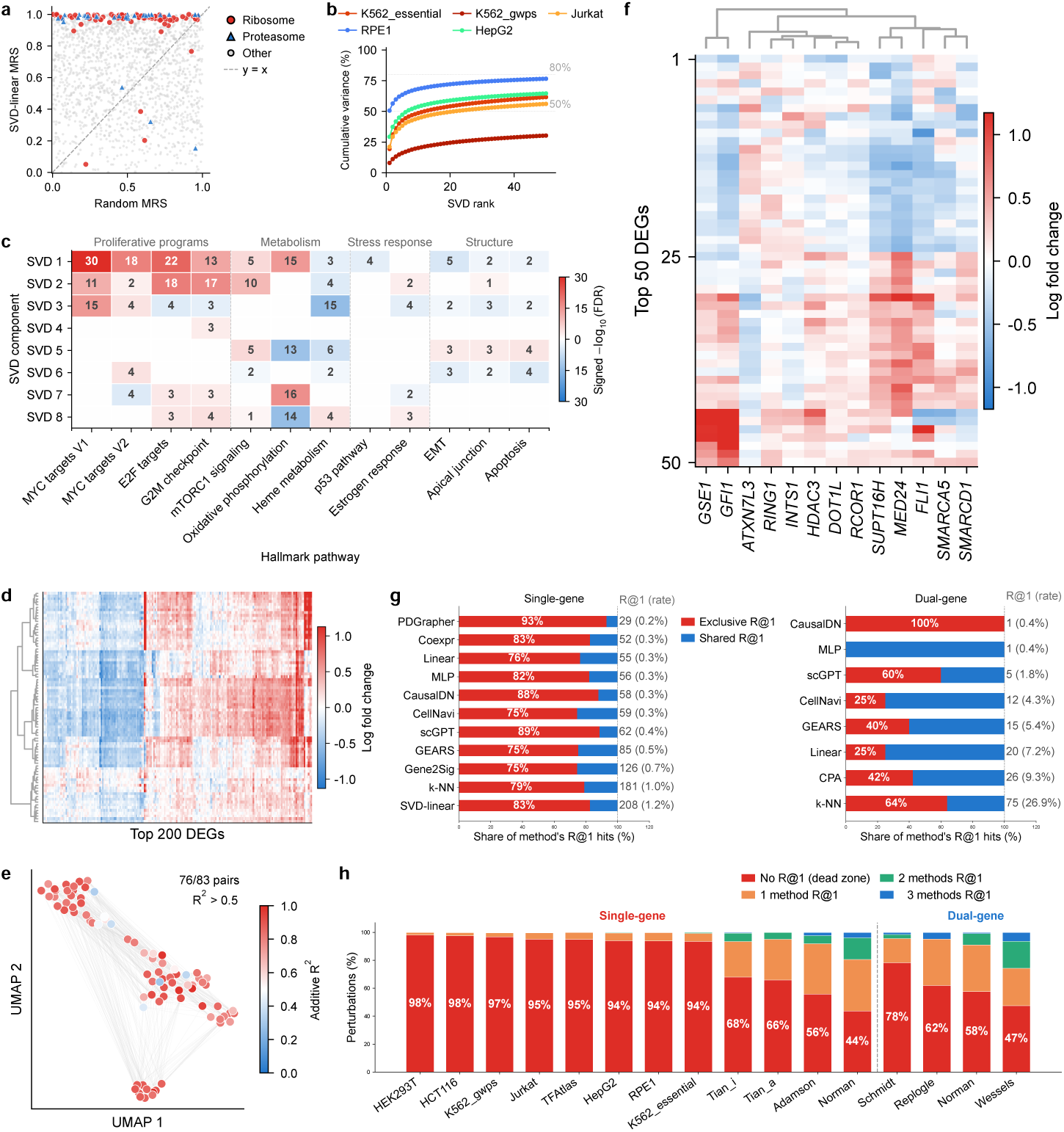
Low-rank structure, additive combinatorial effects, and learnability boundaries in perturbation target identification. **a**, SVD-linear versus Random mean reciprocal score (MRS) per perturbation in K562_essential. Ribosomal protein genes and proteasome subunit genes are highlighted against all other target genes, and the diagonal dashed line marks equal performance (y = x). **b**, Cumulative variance explained by SVD components of the perturbation response matrix (perturbations *×* genes) for five CRISPR interference (CRISPRi) datasets. Dashed lines mark the 50% and 80% thresholds. **c**, Hallmark pathway enrichment of SVD right singular vectors (gene loadings) in K562_essential. Rows are SVD components 1–8, and columns are 13 Hallmark gene sets grouped into four functional categories (Proliferative programs, Metabolism, Stress response and Structure). Cell color shows the signed *−* log_10_(FDR) from a Wilcoxon rank-sum test on gene loadings, and numbers mark the statistic where FDR *<* 0.05 after Benjamini–Hochberg correction. **d**, Hierarchically clustered expression-effect matrix for the 83 dual-gene perturbation pairs in Wessels. Rows are perturbation pairs, columns are the top 200 differentially expressed genes (DEGs), and color encodes log fold change. **e**, UMAP embedding of the 83 dual-gene perturbation pairs in Wessels, colored by additive *R*^2^. Of the 83 pairs, 76 have additive *R*^2^ *>* 0.5. **f**, Additive basis matrix learned by Ridge-kNN in Wessels. Columns are additive basis vectors, each derived from one single-gene perturbation in the training set, rows are the top 50 DEGs, and color encodes log fold change. **g**, Method complementarity in the single-gene and dual-gene benchmarks. Each bar is one method, with bar length encoding the overall R@1 rate and partitioned into hits exclusive to that method and hits shared with at least one other method. Total R@1 hit counts and rates are annotated. **h**, Per-dataset distribution of perturbation retrievability across all 16 benchmark datasets, ordered by decreasing dead-zone fraction within each task. Stacked bars show the fraction of perturbations recovered by zero methods (dead zone), one method, two methods, or three or more methods, with dead-zone percentages annotated. Single-gene and dual-gene datasets are separated by a vertical dashed line.

The dominant singular vectors aligned with recognizable biological programs. In K562_essential, the leading right singular vectors showed significant enrichment for Hallmark gene sets ^27^, including MYC targets, E2F targets and oxidative phosphorylation, among others (Fig. 6c). Because the dominant axes of mathematical variation correspond to known cellular programs, SVD-linear implicitly compares perturbations along biologically interpretable dimensions, explaining its consistent advantage after CEE correction.

The dual-gene results revealed an analogous structural property at the combinatorial level. In Wessels ^25^, additive *R*^2^ exceeded 0.5 for 91.6% of pairs, and the spatial distribution across the pairwise embedding was smooth rather than random (Fig. 6d,e). This geometric regularity provides the statistical basis for Ridge-kNN and explains why combining global additive modeling with local neighborhood adjustment produced robust cross-dataset performance. Single-gene low-rank structure and dual-gene additive structure reflect the same underlying principle. The perturbation response landscape is organized by reusable biological modules, and methods whose inductive bias matches this modularity outperform those that do not.

The additive basis vectors learned by Ridge were also interpretable. In Wessels, the 13 leading basis components organized top differentially expressed genes into coherent expression programs separating activation and repression directions (Fig. 6f), mirroring the Hallmark pathway alignment of SVD singular vectors. This parallel between low-rank projection for single-gene targets and additive decomposition for gene pairs establishes a unified inductive bias principle. In both settings, a method’s prior operates as a linear combination over functional modules, and performance scales with how tightly that prior matches the modular geometry of the perturbation response landscape.

The benchmark also revealed substantial complementarity among structured methods. Most correctly identified perturbations were recovered by only one method (Fig. 6g), indicating that current approaches capture partially non-overlapping subsets of the recoverable perturbation space. Ensemble strategies combining low-rank, neighborhood and gene-feature signals could therefore increase total coverage without additional training data.

The dead-zone analysis quantified this limitation across the full benchmark. For each of the 16 datasets, we classified every test perturbation by how many fair methods achieved R@1 success on it. In the majority of datasets, the fraction recovered by no method exceeded 50%, with several genome-wide datasets above 90% (Fig. 6h). The dead zone is highest in unstructured genome-wide screens such as K562_gwps (*∼*90%). Pathway-focused panels such as Norman and Adamson, with dead-zone fractions near 50%, demonstrate that concentrating perturbations within coherent biological programs substantially increases fair learnability. The approximately 8% of pairs in Wessels with additive *R*^2^ below 0.5 likely concentrate on biologically critical epistatic interactions. Targeted identification of these cases, rather than improving overall additive performance, represents the more precise research objective. Achieving it will require combinatorial screens that systematically sample gene pairs within and across functional modules at sufficient scale.

## 3 Discussion

We introduced CEE to evaluate perturbation target identification under leakage-free conditions. Training leakage arises when test-perturbation cells are observed during model fitting; evaluation leakage arises when target-gene self-effects remain in the query signature and enable direct retrieval without modeling downstream biology. Applied to 16 single-cell perturbation datasets, the framework changed the apparent method hierarchy. Among the 12 benchmarked approaches, response-imputation methods such as SVD-linear and k-NN, and gene-feature methods such as Gene2Sig, retained the strongest CEE signal, whereas standard and upper-bound evaluation settings overstated the performance of self-effect-driven or train-exposed methods. This reversal matters for cell-state design, where the objective is not to rank a held-out signature but to recover an actionable intervention from downstream biology. Two structural features of the current landscape bound how far the task can presently be pushed. First, models built natively for the inverse direction remain few, comprising essentially CausalDN, PDGrapher and CellNavi ^9–11^, so that most comparisons still rely on repurposed forward predictors or statistical baselines. Second, combinatorial perturbation atlases remain scarce and sample only a small fraction of possible gene pairs ^3,23–25,28^. The low absolute performance under CEE evaluation therefore marks an early methodological stage, not a limit on the task itself.

Our findings are consistent with a broader pattern emerging across perturbation benchmarking. Recent work on forward prediction has shown that simple baselines frequently match or outperform deep learning models when evaluation protocols control for systematic variation ^16,17,29^. CEE extends this insight to the inverse direction and isolates a mechanism specific to target identification: the self-effect of the perturbed gene can be read directly from the query signature, an artefact absent from forward benchmarks but dominant in conventional inverse evaluation. Together, these results suggest that single-cell perturbation modeling has systematically overestimated the capability of complex models, in both directions, through protocols that inadvertently reward artefact exploitation rather than biological generalization. Yet the inverse direction is where much of the practical demand lies. Target discovery, mechanism-of-action analysis, drug repurposing and rational cell-state engineering all begin from a desired transcriptional or phenotypic state and ask which intervention reaches it ^4–8,30^. The field has nonetheless concentrated overwhelmingly on forward response prediction, leaving the inverse task served largely by forward models repurposed after the fact. We therefore see a clear need for methods designed from the outset for target identification, optimized against downstream response rather than reconstruction fidelity, and benchmarked under leakage-free protocols such as CEE.

How such methods should be built follows from the structure that survives CEE correction. The recoverable single-gene signal is low-rank and aligns with recognizable functional programs. This argues for inductive biases that operate explicitly over functional modules, such as low-rank projections or signature bases, and for training objectives defined directly in the downstream response space rather than on the raw query profile. It also argues against priors that mismatch this geometry. A protein– protein interaction prior actively impaired functional recovery relative to data-driven signatures, indicating that network topology is not interchangeable with functional similarity. The dual-gene setting is governed instead by additive decomposability. Progress there therefore hinges less on raising overall additive accuracy than on the minority of non-additive, epistatic pairs that additive models cannot capture by construction, and on combinatorial screens that sample gene pairs systematically within and across functional modules. Our learnability analysis further offers a concrete experimental prescription. At fixed sequencing budget, pathway-focused perturbation panels yielded substantially higher fair learnability than unstructured genome-wide screens, so concentrating perturbations within coherent biological programs is a more productive design choice than scale alone. In the near term, the limited overlap among the perturbations recovered by different structured methods indicates that ensembling low-rank, neighborhood and gene-feature signals could broaden coverage without any new data. The deeper lesson of CEE, however, is one about evaluation itself. Self-effect leakage is not a peripheral artefact but a structural misalignment between what benchmarks have measured and what the task is for, allowing a model to top a leaderboard by reading the cause off its own effect rather than reasoning from effect back to cause. Removing this shortcut is what makes the inverse problem honest. We therefore argue that target identification methods should ultimately be judged by their biological utility, namely whether the interventions they nominate actually move cells toward desired states, an assessment that emerging high-content readouts such as optical pooled and spatial functional genomics screens are now beginning to make directly testable.

## 4 Methods

### Gene expression perturbation datasets

We assembled 16 public single-cell perturbation datasets for reverse perturbation inference. The single-gene benchmark included 12 datasets generated by pooled single-cell CRISPR and Perturb-seq screening designs ^1,31^: Adamson2016^18^, HepG2 and Jurkat ^19^, K562_essential, K562_gwps and RPE1^3^, Tian2021_CRISPRa and Tian2021_CRISPRi ^20^, TFAtlas ^21^, HCT116 and HEK293T ^22^, and Norman_K562_single ^23^. These datasets covered CRISPRi, CRISPRa and ORF perturbations across cell lines and perturbation-panel designs. The dual-gene benchmark included four datasets: Norman_K562_dual ^23^, Schmidt ^24^, Wessels ^25^ and Replogle2020^26^. These datasets covered CRISPRa, Cas13 RNA-targeting knockdown and CRISPRi assays, including combinatorial perturbation designs used for genetic-interaction measurements ^32^.

#### Dataset standardization

We converted all datasets to a common AnnData schema using the Scanpy single-cell analysis framework ^33^. Perturbation labels were stored as target_gene, gene-symbol labels as target_gene_symbol, and non-targeting or control cells as nontargeting. For datasets distributed as count matrices, we normalized each cell to a total count of 10^4^ and log-transformed the result. For datasets distributed as log-normalized expression matrices, we retained the supplied log-transformed values. Norman single-gene and dual-gene subsets were derived from the same processed source dataset by separating single-target conditions from two-gene combinations.

### Data preprocessing

#### Gene selection

For each dataset, we selected a benchmark gene space of up to 5,000 highly variable genes (HVGs), a standard representation in single-cell integration workflows ^34^. We computed gene-level variance in log-normalized expression and selected the detected genes with the highest variance after requiring expression in at least three cells. Perturbation target genes were forced into the benchmark gene space whenever they were present in the raw gene annotation, even if they were not among the highest-variance genes. For log-normalized datasets with no more than 5,000 genes after standardization, we retained all detected genes rather than applying an additional HVG filter.

#### Perturbation filtering

We removed perturbation labels represented by fewer than five cells. For genome-scale or very large datasets, we limited the benchmark to the most frequently observed perturbations while retaining all control cells. The cap was 3,000 perturbations for K562_gwps, HCT116 and HEK293T. We also applied a perturbation-direction filter based on the expected target-gene log fold-change direction. CRISPRi and Cas13 perturbations were required to decrease target-gene expression, whereas CRISPRa and ORF perturbations were required to increase it.

### Cause-excluded effect evaluation framework

#### Reverse perturbation inference

Let *G* denote the number of genes in the benchmark expression space. For a perturbation query *q*, the differential expression profile between perturbed and control cells is defined as the mean perturbed expression minus the mean control expression across *G* genes. In expression-signature target-identification and connectivity-mapping analyses, perturbation profiles are compared with candidate-specific signatures ^4,5,7,8,30^. We evaluated all retrieval-style methods by comparing query profiles with candidate-specific predicted or observed profiles in the same gene space. Formally,

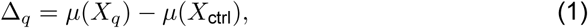

where *µ*(*·*) denotes the cell-mean operator and Δ*_q_ ∈* R*^G^*. Reverse perturbation inference ranks a candidate target set *C* and asks whether the true target gene, or the true target-gene pair in the dual-gene setting, receives a high rank.

#### Profile-wise self-effect exclusion

We used CEE to exclude direct target-gene self-effects from each query while preserving downstream expression consequences. This exclusion replaced the target-gene coordinates with a fold-specific non-self background rather than dropping them from the gene space. The CEE evaluation setting corresponds to the CEE protocol reported in the Results and figure legends. For fold *f*, gene *g* and the train-only perturbation sample set *T_f_*, we estimated the non-self background expression as

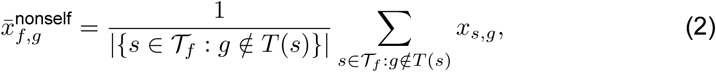

where *T* (*s*) is the set of targeted genes in sample *s*. For a query *q*, each target-gene coordinate *g ∈ T* (*q*) was replaced with this fold-specific non-self background before subtracting the control mean:

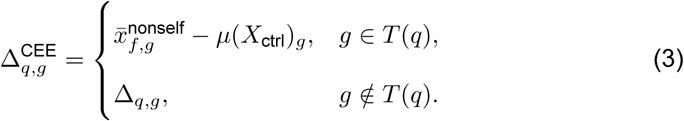

For single-gene queries, one coordinate was replaced. For dual-gene queries, both target coordinates were replaced independently using the fold-specific background for each gene. We applied the same replacement to candidate signatures generated by signature-based methods and to the training inputs of classifiers and fine-tuned external models. The same preprocessing rule was used at both training and evaluation stages. Similarity scores were computed as cosine similarity in the full benchmark gene space R*^G^*, rather than in a reduced gene subset.

#### Cell-level self-effect exclusion

For models that consume single-cell expression matrices directly, we applied column-level resampling to the target gene before model input construction. This procedure was used for CausalDN, which derives input features from per-gene mean, variance and cross-gene correlation computed across a batch of cells. For target gene *g*, the target column in both the perturbed-cell and control-cell matrices was replaced with independent draws from held-out empirical distributions:

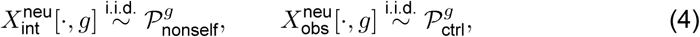

Where ρ*^g^*_nondself_ is the empirical distribution of gene *g* expression across all fold train-only perturbed cells whose perturbation target is not *g*, and ρ*^g^*_ctrl_ is the empirical distribution of gene *g* in control cells. All *G −* 1 other columns remained unchanged. The resampled matrices were then passed to the corresponding cell-level model pipeline.

#### Evaluation protocols

We used three evaluation settings. In the CEE evaluation setting, train and test perturbations were separated at the perturbation level, and queries were evaluated after profile-wise self-effect exclusion. In the standard evaluation setting, train and test perturbations were also separated at the perturbation level, but the raw query self-effect was retained. In the upper-bound evaluation setting, test perturbation cells could be observed during model fitting or method construction. The fair settings excluded held-out perturbation information from model construction and feature engineering ^15^.

#### Self-effect measurements

We quantified the target-gene self-effect in two complementary ways. For energy-based analyses, we used the self-effect energy fraction

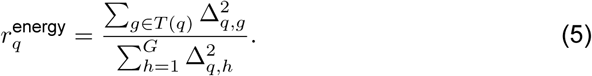

For differential gene expression (DGE) dominance analyses, we used the self-effect dominance ratio

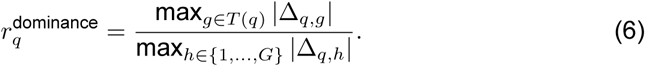

We classified a single-gene query as DGE-trivial when the true target gene was the largest-magnitude component of the raw differential expression profile.

### Cross-validation protocol

#### Fold construction

We used perturbation-level cross-validation for both single-gene and dual-gene benchmarks, as in recent evaluations of generalizable perturbation response prediction ^16,17^. Each dataset was split into five folds, with test perturbations held out from model fitting in each fold. For datasets with fewer than 100 perturbations (Adamson2016, Tian2021_CRISPRa, Schmidt and Replogle2020), three-fold cross-validation was used instead to maintain an adequate number of test perturbations per fold and ensure stable performance estimates. The serialized fold metadata under processed_data/folds/ was used as the single source of truth for train-only, validation and test perturbation identities. Methods with validation-based training control used the validation perturbations from the same fold. Nonparametric and closed-form methods were fit with the fold-specific training perturbations.

#### Fold-specific non-self background

We estimated the non-self background independently within each fold using only train-only perturbation samples. The resulting vector was cached as train_only_nonself_bg.npy in the corresponding fold directory. Validation and test perturbations were not used to estimate this vector.

### Single-gene reverse perturbation benchmark

#### Native reverse methods

We evaluated CausalDN ^9^, CellNavi ^11^ and PDGrapher ^10^ as native reverse or target-identification models in the single-gene benchmark. In the CEE evaluation setting, each method used fold-specific predictions or inference outputs that excluded test perturbations from the fitted model state. PDGrapher used the CEE-compatible inference output in which target-gene self-effect information was masked. We also evaluated no-mask and upper-bound variants for leakage analyses, but these variants were not included in the main CEE ranking.

#### Forward prediction models

We repurposed GEARS ^12^ and scGPT ^13^ for retrieval in the single-gene benchmark. Related single-cell perturbation response models include scGen, PRESCRIBE and scPRAM ^35–37^. For each fold, a forward model predicted a differential expression profile for each candidate target gene. We excluded the candidate target coordinate and ranked candidates by cosine similarity between the CEE-corrected query profile and the predicted candidate profiles.

#### Response imputation methods

SVD-linear constructed a perturbation-by-gene response matrix from training perturbations, performed truncated singular value decomposition using SciPy ^38^ and learned a linear map from gene-coordinate loadings to perturbation-coordinate loadings. Candidate signatures were reconstructed in the low-rank response space and evaluated by full-space cosine similarity after target-coordinate exclusion. The SVD rank was selected per dataset using mean reciprocal rank (MRR)-optimized five-fold ablations, with ranks in *{*20, 30, 50*}* depending on the dataset. The k-NN method estimated each candidate gene’s response signature as a weighted average of training perturbation signatures from genes with similar coexpression profiles. The value of *K* was selected per dataset using the same cross-validation protocol.

#### Gene feature methods

Gene2Sig trained an MLP regressor from gene coexpression features to CEE-corrected perturbation signatures, using scikit-learn estimators for tabular regression and model fitting ^39^. Coexpr used the fold-specific coexpression matrix directly as a candidate gene feature representation and scored candidates by full-space cosine similarity after target-coordinate exclusion.

#### Reverse classifier methods

Linear and MLP classifiers directly mapped a differential expression profile to a multilabel target-gene score vector. The linear classifier used one fully connected layer, and the MLP used two hidden layers with 512 and 256 units, following standard feedforward-network parameterizations ^40^. Both models were implemented in PyTorch ^41^ and trained with the Adam optimizer ^42^ within each fold with CEE-corrected training and inference inputs in the CEE evaluation setting.

#### Upper-bound references

DGE ranked genes by the magnitude of their observed differential expression and was treated as an upper-bound reference that does not participate in fair ranking. CPA ^14^ was included as a fair-ranking participant in the single-gene benchmark, evaluated under the CEE unseen-perturbation split with self-effect exclusion applied at training and inference.

### Dual-gene reverse perturbation benchmark

#### External methods

We evaluated CellNavi, CausalDN, PDGrapher, GEARS, CPA and scGPT in the dual-gene benchmark. In the CEE evaluation setting, test gene pairs were excluded from training. Single-gene perturbation cells were included when required by a method, whereas held-out pair responses were excluded. GEARS, CPA and scGPT produced candidate pair expression-effect signatures that were scored by full-space cosine similarity after pair-target exclusion. CellNavi and CausalDN produced direct candidate scores, whereas PDGrapher compared predicted pair-level outputs with candidate expression-effect signatures.

#### Additive retrieval methods

Ridge modeled each pair response as an additive combination of latent single-gene effects. For training pairs (*a, b*), we constructed a design matrix *A* with one indicator for each gene in the pair and fit

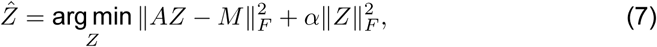

where *M* is the matrix of CEE-corrected pair response signatures. We used *α* = 0.5 for Ridge. Ridge-kNN first fit the same additive model with *α* = 0.3, then blended the additive prediction with a local weighted average of the five nearest training-pair signatures. The final signature was 0.6 times the additive prediction plus 0.4 times the local k-NN prediction.

#### Simple baselines and upper-bound reference

The dual-gene k-NN baseline estimated query-specific single-gene effects from train ing pairs that contained the relevant gene. Candidate pairs were scored by cosine similarity between the query and the sum of the two estimated effects. Dual-gene Linear and MLP classifiers used the same model classes as the single-gene classifiers, with pair scores computed as the sum of the two predicted gene logits. DGE-pair was retained as an upper-bound reference that used the observed expression changes of candidate pair genes and was not included in the fair ranking.

### Performance metrics

#### Ranking task definition

For each query, every method returned a score *s*(*c*) for each candidate *c ∈ C*. Candidates were ranked in decreasing score order. If tied scores occurred, we used the average rank among tied candidates. A single-gene query was correct at rank *k* if its true target gene appeared within the first *k* candidates. A dual-gene query was correct at rank *k* if the exact unordered target pair appeared within the first *k* candidates.

#### Ranking metrics

We reported recall at *k*, MRR, mean reciprocal score (MRS), normalized discounted cumulative gain at 10 (NDCG@10) ^43^ and AUROC. For a query with target rank *r_q_* among *N_q_* candidates,

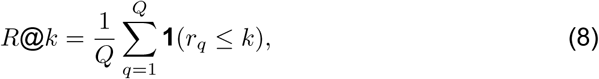

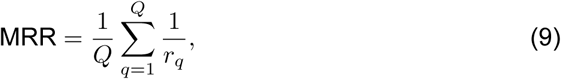

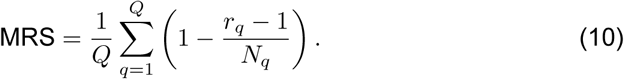

For a single relevant target, NDCG@10 was 1/ log_2_(*r_q_*+ 1) when *r_q_ ≤* 10 and zero otherwise. AUROC had the closed-form value

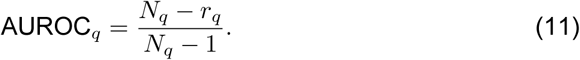

For dual-gene queries, we also reported Partial@1, Partial@5 and Partial@10, where a partial hit occurred if any candidate in the top *k* shared at least one gene with the true pair.

#### Composite rank

To summarize performance across datasets, we ranked methods within each dataset using the primary metrics reported in the corresponding figure and averaged the resulting within-dataset ranks. This composite rank was used only as a cross-dataset summary.

### Leakage analysis

#### Self-effect leakage analysis

We analyzed evaluation leakage by comparing DGE ranks with self-effect measurements and by perturbing the target-gene coordinate directly. In the controlled injection experiment, we multiplied the target-gene coordinate of a query by a scale factor *λ* and recomputed DGE ranks across values of *λ*.

#### Training leakage protocol

We quantified training leakage by comparing upper-bound variants with CEE or standard evaluation variants under matched fold definitions. CausalDN, CellNavi and PDGrapher were evaluated in CEE-compatible, no-mask and upper-bound forms where available. CEE-compatible variants excluded test perturbations and applied the appropriate self-effect masking or profile-wise self-effect exclusion. No-mask variants excluded test perturbations from training but retained the raw query self-effect. Upper-bound variants allowed test perturbation information during model fitting or fine-tuning.

## Acknowledgements

This work was supported by the Strategic Priority Research Program of the Chinese Academy of Sciences (XDB1350302), the National Key Research and Development Program of China (2024YFF0729200), the National Natural Science Foundation of China (32341013), the National Key Research and Development Program of China (2023YFA1802003), the Strategic Priority Research Program of the Chinese Academy of Sciences (XDA0460305, XDA0510401, and XDC0200102), the Initiative Scientific Research Program, Institute of Zoology, Chinese Academy of Sciences (2024IOZ0101 and 2023IOZ0102), and the Open Project Fund of the State Key Laboratory of Organ Regeneration and Reconstruction, Chinese Academy of Sciences (2026KF11).

## Author contributions

J.H., X.Q. and X.L. conceived and designed the study. J.H. developed the CEE framework and the evaluation methodology, with methodological input from J.Y. J.H. implemented the software, performed the formal analysis and curated the data. J.H., L.T. and N.L. contributed to software development, validation and visualization. J.Y. contributed to methodology and validation. X.Q. and X.L. supervised the project and provided resources and project administration. X.L. acquired funding. J.H. wrote the original draft. X.Q. and X.L. reviewed and edited the manuscript. All authors read and approved the final manuscript.

## Data availability

All 16 single-cell perturbation datasets analyzed in this study are publicly available. Adamson2016 is available at GEO under accession GSE90546. Norman2019 (single-gene and dual-gene subsets) is available at GSE133344. Replogle2022 (K562_essential, K562_gwps and RPE1) is available via SRA accession PRJNA831566 and processed files at Figshare (https://doi.org/10.25452/figshare.plus.20029387). Nadig2024 (HepG2 and Jurkat) is available at GSE264667. Tian2021 (CRISPRa and CRISPRi) is available at GSE152988. TFAtlas (Joung2023) is available at GSE216481. Schmidt2022 is available at GSE190604 (companion series GSE174292 from the same study); processed files are also accessible via scPerturb Zenodo (https://doi.org/10.5281/zenodo.7041848). Wessels2023 is available at GSE213957 and via scPerturb Zenodo (https://doi.org/10.5281/zenodo.7041848). Replogle2020 is available at GSE146194. The HCT116 and HEK293T datasets (Huang2025) are available from HuggingFace at https://huggingface.co/Xaira-Therapeutics/X-Atlas-Orion. Processed benchmark datasets and fold definitions used in this study are available at https://github.com/Hjxzuibang/CEE.

## Code availability

Source code for the CEE evaluation framework, all baseline evaluations and figure reproduction scripts is available at https://github.com/Hjxzuibang/CEE (archived at Zenodo: https://doi.org/10.5281/zenodo.20850789) under the Apache License 2.0. The framework provides a Python API and a command-line interface, together with end-to-end shell scripts for preprocessing, evaluation and figure generation. Full installation and usage instructions, including an example notebook, are provided in the repository documentation.

